# Long non-coding RNA GAS5 acts as proliferation “brakes” in CD133+ cells responsible for tumor recurrence

**DOI:** 10.1101/670968

**Authors:** Nikita S Sharma, Prisca Gnamlin, Brittany Durden, Vineet K Gupta, Kousik Kesh, Vanessa T Garrido, Roey Hadad, Vikas Dudeja, Ashok Saluja, Sulagna Banerjee

**Author notes:** Contributed equally (Nikita S Sharma, Prisca Gnamlin). Corresponding Author: Address of Correspondence: 1551 NW 10^th^ Ave, Biomedical Research Building, Suite 508 Miami, FL 33136.

## Abstract

Presence of quiescent, therapy evasive population often described as cancer stem cells (CSC) or tumor initiating cells (TIC) is often attributed to extreme metastasis and tumor recurrence. This population is typically enriched in a tumor as a result of microenvironment or chemotherapy induced stress. The TIC population adapts to this stress by turning on cell cycle arrest programs that is a “fail-safe” mechanism to prevent expansion of malignant cells to prevent further injury. Upon removal of the “stress” conditions, these cells restart their cell cycle and regain their proliferative nature thereby resulting in tumor relapse. Growth Arrest Specific 5 (GAS5) is a long-noncoding RNA that plays a vital role in this process. In pancreatic cancer, CD133+ population is a typical representation of the TIC population that is responsible for tumor relapse. In this study, we show for the first time that emergence of CD133+ population coincides with upregulation of GAS5, that reprograms the cell cycle to slow proliferation by inhibiting GR mediated cell cycle control. The CD133+ population further routed metabolites like glucose to shunt pathways like pentose phosphate pathway, that were predominantly biosynthetic in spite of being quiescent in nature but did not use it immediately for nucleic acid synthesis. Upon inhibiting GAS5, these cells were released from their growth arrest and restarted the nucleic acid synthesis and proliferation. Our study thus showed that GAS5 acts as a molecular switch for regulating quiescence and growth arrest in CD133+ population, that is responsible for aggressive biology of pancreatic tumors.

## Introduction

Aggressiveness of a tumor has been correlated with the presence of a population of slow-cycling, treatment refractory and extremely metastatic cells. Accumulating evidence shows that this population is typically enriched in a tumor in response to microenvironmental and/or chemotherapy induced stress. Recent research has attributed this enrichment to senescence associated “stemness” ^22^. These studies have shown that under chemotherapeutic or microenvironmental stress like hypoxia or nutrient deprivation, a population of cells specifically respond to the induced stress by triggering a cell cycle arrest program that prevents further expansion of the malignant cells. This is considered to be a failsafe mechanism by the tumor to prevent further “injury”. Upon removal of the stress, this population promptly regains its proliferative nature, thereby leading to relapse and recurrence of the tumor.

Pancreatic adenocarcinoma is notorious for its resistance to therapy, metastasis and high rate of recurrence (www.cancer.gov). Studies from our laboratory show that a CD133+ population is associated with the aggressive biology of pancreatic adenocarcinoma^2^. While they are probably not a population that is responsible for the origin of pancreatic tumors, our previously published study definitely show that they are responsible for therapeutic resistance, tumor initiation at very low dilution as well as extreme metastasis^2, 24, 26^. Our studies further show that this population is enriched upon nutritional deprivation, low dose chemotherapy as well as presence of hypoxia^21, 25, 26^. We and others have shown that CD133+ population are generally slow-cycling or quiescent^2, 6, 36^. This indicates that the cell cycle plays an active role in maintenance of this population in a quiescent and slow cycling state.

Growth Arrest Specific 5 or GAS5, is a long non-coding RNA regulates cell cycle in a number of mammalian systems including several cancers ^7, 15, 16, 23^. It also mediates cell proliferation by regulating CDK6 activity ^17^. Studies have also shown that GAS5 forms a positive feedback network with a number of genes involved in self-renewal like Sox2/Oct4, making this long non-coding RNA (LncRNA) a critical player in induction and maintenance of the “stemness” state in a tumor ^34^. GAS5 is further involved in regulation of human embryonic stem cell self-renewal by maintaining NODAL signaling^38^. Mechanistically, the effect of GAS5 on cell cycle is regulated by its interaction with the glucocorticoid receptor (GR)^11^. GRs are nuclear receptor proteins that control cell proliferation via their effect on cell cycle ^30^. GAS5 interacts with the activated GR preventing its association with the glucocorticoid response element (GRE) and consequently suppressing the transcription of target genes ^19^.

Studies from our laboratory have shown that the CD133+ population of cells is metabolically reprogrammed to be more dependent on glycolysis and has very low dependence on oxidative phosphorylation. Further, our studies have shown that this altered metabolic state promotes a “survival advantage” in this population by minimizing ROS accumulation^26^. Interestingly, while increased aerobic glycolysis is typically thought to be associated with proliferation, recent studies show that this metabolic activity may also be associated with other cellular functions as well ^10^. Increased glucose uptake and metabolism is thus not necessarily required for robust growth of cells ^12, 35^. Literature also shows that glycolysis can be regulated by glucocorticoid receptor (GR) ^5, 18^, and this can further affect cell cycle.

While CD133+ cells have been shown to be aggressive and therapy resistant, the molecular circuitry behind this still remains an enigma. It is also not clear why this population emerges in pancreatic tumors in response to microenvironmental as well as chemotherapeutic stress. In this study, we show for the first time that emergence of CD133+ population coincides with upregulation of GAS5, that reprograms the cell cycle to slow proliferation by inhibiting GR mediated cell cycle control. The CD133+ population further routed metabolites like glucose to the pentose phosphate pathway, that were predominantly biosynthetic in spite of being quiescent in nature but did not use immediately for nucleic acid synthesis. Upon inhibiting GAS5, these cells were released from their growth arrest and restarted the nucleic acid synthesis and proliferation. Our study thus showed that GAS5 acts as a molecular rheostat for regulating quiescence and growth arrest in CD133+ population, that is responsible for aggressive biology of pancreatic tumors.

## Materials and Methods

### 1. Cells and reagents

Human cDNA CD133 expression plasmid (EX-Z0396-M02) and empty vector plasmid (EX-NEG-M02) were obtained from GeneCopoeia. MIA PaCa-2, SU86.86 cells were obtained from ATCC and maintained as recommended. Stable MIA-derivatives (CD133Hi cells) were maintained in DMEM (Hyclone) containing 10% fetal bovine serum. S2-VP10 cells were cultured in RPMI 1640 (Hyclone) supplemented with 10% fetal bovine serum. Stable clones were selected and maintained in Geneticin (Invitrogen).

### 2. Isolation and metabolic labeling of CD133+ tumor initiating cells from tumors

The CD133^+^ population was separated from the mouse progenitor cells and other CD133^−^ cells using MACS separation (Miltenyi Biotech) following manufacturers protocol. Single cell suspension was generated from tumors in KPC mice according to Li *et al* ^13^. Non-epithelial progenitor cells were removed using anti-CD31-Biotin (BD Biosciences) and anti-CD45 Biotin (BD Bioscience) using MACS technique. The flowthrough free from the mouse progenitor cells was bound to anti-mouse CD133-Microbeads for 10 min on ice and positively purified for CD133^+^ cells by MACS. The purity of separation was tested for each batch by performing a FACS analysis using Anti-CD133-PE antibody AC141 (Miltenyi Biotech). The separated populations were used for RNA, Protein and FACS analysis. Cells growing in culture were scraped gently into centrifuge tube and washed once in Wash Buffer (PBS, 0.5% BSA, 2mM EDTA) before binding to Anti-mouse CD133 microbeads and proceeding as described above.

For labeling with ^13^C glucose, isolated CD133^+^ cells were incubated in glucose free DMEM for 30 min. Following incubation, the cells were transferred to a growth medium (DMEM) with ^13^C glucose (final concentration 25mM) and incubated further for 30min. Labeled cells were flash frozen and sent to University of Michigan Regional Comprehensive Metabolomic Resource Core for flux analysis. CD133^−^ cells were processed in parallel with ^12^C glucose (final concentration 25mM) added to DMEM.

### 3. Cell Cycle analysis

Pancreatic cancer cells were plated in a 6-well plate and grown till 70-80% confluent. Following this, cells were harvested by trypsinization, washed and fixed in 70% cold ethanol, taking care to add ethanol dropwise while gently vortexing the cells. Cells were fixed at 4°C for 30 min, washed and treated with RNAse. A 100 µl aliquot of this suspension was taken and mixed with 100 µl PI (Sigma Aldrich) reagent, incubated in dark for 30 minutes and then acquired on BD FACS Canto II. Cell cycle analysis was done by measuring total DNA content of the cells. For analysis in CD133+ vs CD133- cells, SU86.86 cells and/or KPC001 cells were sorted by staining with anti-CD133-PE in BD-FACS Aria. Sorted cells were grown for 2 weeks in growth media supplemented with 10% FBS. Once 70% confluent, analysis was done as described above.

For Pyronin Y staining, cells were grown or sorted as described above. For staining, cells were resuspended in 1ml cell culture medium containing 10µg/ml Hoechst 33342 and incubated at 37°C for 45 min. 5µl of 100µg/ml Pyronin Y (Sigma Aldrich) was added to the cells and incubated for another 15 minutes. Cells were analyzed in BD FACS CantoII.

### 4. Glucocorticoid Receptor Activity Assay

CD133Hi and controls were seeded in a 24 well plate. Cells were transfected with the Cignal Reporter Plasmids for GRE using a dual luciferase assay system (Qiagen). Appropriate treatment (siGAS5, dexamethasone, etc) was done for 24 hours. The dual luciferase kit (Promega) was used to measure activity using a luminometer. Each sample was treated in duplicate for each plasmid (the negative reporter the GRE reporter).

### 5. G6PD Assay

G6PD activity assay was performed on CD133hi and CD133lo cells using the G6PD activity assay kit (Sigma-Aldrich) according to manufacturer’s protocol.

### 6. BrDU incorporation assay

For BrDU incorporation and nucleic acid synthesis in CD133hi and CD133lo cells, cells were treated with 10ug/ml BrDU (BD Biosciences) following Sox2 or GAS5 silencing. Staining for BrDU was done according to manufacturer’s instruction.

### 7. Proliferation Assay

For proliferation, MTT based assay was used. Briefly, cells were plated in 100ul growth medium in a 96-well plate and appropriate treatment was done for 24h,48h or 72h. Following treatment, 10ul CCK-8 reagent was added to each well and incubated at 37°C for 1h. The plate was read at 450nm in a Molecular Devices spectrophotometer. Data was normalized to time 0 to get proliferation rate.

### 8. Fluorescent In Situ Hybridization

For FISH with GAS5 probe (Stellaris), FFPE slides were deparaffinized in 100% xylene for 10 minutes and additionally in fresh 100% xylene for 5 minutes. Slides were then dipped in 100% EtOH for 10 min, again in fresh 100% EtOH for 10 minutes, and finally 70% EtOH for at least one hour. Slides were incubated in 1x PBS for 5 min followed by 20 minute incubation with prewarmed proteinase k solution (10ug/mL proteinase k in 1X PBS warmed to 37°C).

Hybridization was performed by immersing slides in wash buffer A (30 mL wash buffer A from Stellaris kit, 105mL dH2O, 15mL deionized formamide) for 5 min. Slides were removed from wash buffer A and excess was wiped off. 200uL hybridization buffer (2uL Gas5 probe stock + 200uL hybridization buffer) were added to each slide and a cover slip was applied to the top. Slides were incubated in the dark overnight at 37°C for no more than 16 hours.

Slides were immersed in wash buffer A to allow cover slip to detach and then incubated in the dark at 37°C for 30 min. ProLong Gold anti-fade reagent with DAPI was added to each slide and a new cover slip applied. Slides were stored in 4°C in the dark and imaged with Leica fluorescent microscope.

### 9. Statistics

Values are expressed as mean ± SEM. *In vitro* were performed a minimum of three times and significance between two samples was determined using the Student’s paired t-test. Values were considered statistically significant when p<0.05.

## Results

### CD133^Hi^ cells are non-proliferative and arrested in G0/G1 phase

It is well known that "stem" cells are non-proliferative and quiescent. In our previous studies, we determined that pancreatic cancer cell lines like S2VP10, SU86.86 and KPC001 had higher number of CD133+ cells compared to MIAPACA2 and PanC01^2^. We called the cell lines with increased CD133+ population, CD133^Hi^ while the cells with less CD133+ cells CD133^Lo^. Additionally, we also overexpressed CD133 constitutively in MIA-PACA2 cell line (that have negligible CD133+ population) to generate a MIA-CD133^Hi^ cell line^24^. To study if all cell lines with increased CD133 population were non-proliferative, we next tested the proliferation of cells with varied CD133 expression level. Our results showed that cells with increased CD133+ population were less proliferative compared to cell lines with less CD133+ population (Figure 1A, B). This was similar to what was observed in isolated CD133+ cells from KPC tumors as well (Supplementary Figure 1A). To study if this decreased proliferation was due to an arrest in cell cycle, we isolated CD133+ cells from KPC001 and SU86.86 cells and performed a cell cycle analysis by propidium iodide staining. Our studies showed that CD133+ cells (from both these cell lines) were arrested in G0/G1 phase with a significant increase in the G0 Phase cells, while the CD133- cells progressed to G2/M phase regularly (Figure 1C,D, Supplementary Figure 1B). This was further confirmed by a Hoechst 33342/Pyronin Y staining. Generally, resting/quiescent cells at G0 phase have lower levels of RNA compared with proliferating interphase cells (G1-S-G2-M phase). Pyronin Y intercalates both double stranded DNA and double stranded RNA. In the presence of DNA-chelating fluorescent dye such as Hoechst 33342, interactions of Pyronin Y and DNA complex are disrupted and Pyronin Y mainly stains RNA ^31^, allowing the quantification of RNA amount in a single cell level. Thus, resting cells can be identified by lower Pyronin Y for the same amount of DNA. Our results showed that CD133+ cells had an increase in low Pyronin stained population compared to CD133- cells in cells isolated from SU86.86 as well as KPC001 cell lines (Figure 1 E, Supplementary Figure 1C). To further understand the mechanisms of arrest we next analyzed the cell cycle regulator genes. Among these, cyclin dependent kinase 4/6 (CDK4/6) is important for G1 phase progression and is regulated by cyclin D. Cyclin D forms a complex with CDK4 to phosphorylate and inhibit retinoblastoma, Rb and regulates the G1/S transition. Our results showed that in CD133hi cells had increased expression of CDK4, CDK6 and Cyclin D2 (Figure 1F, G, Supplementary Figure 1D) and increased phosphorylation of Rb (Figure 1G). This indicates that in CD133+ cells, CDK4 expression and activity was higher than in CD133- cells leading to a G0/G1 arrest.

**Figure 1.**
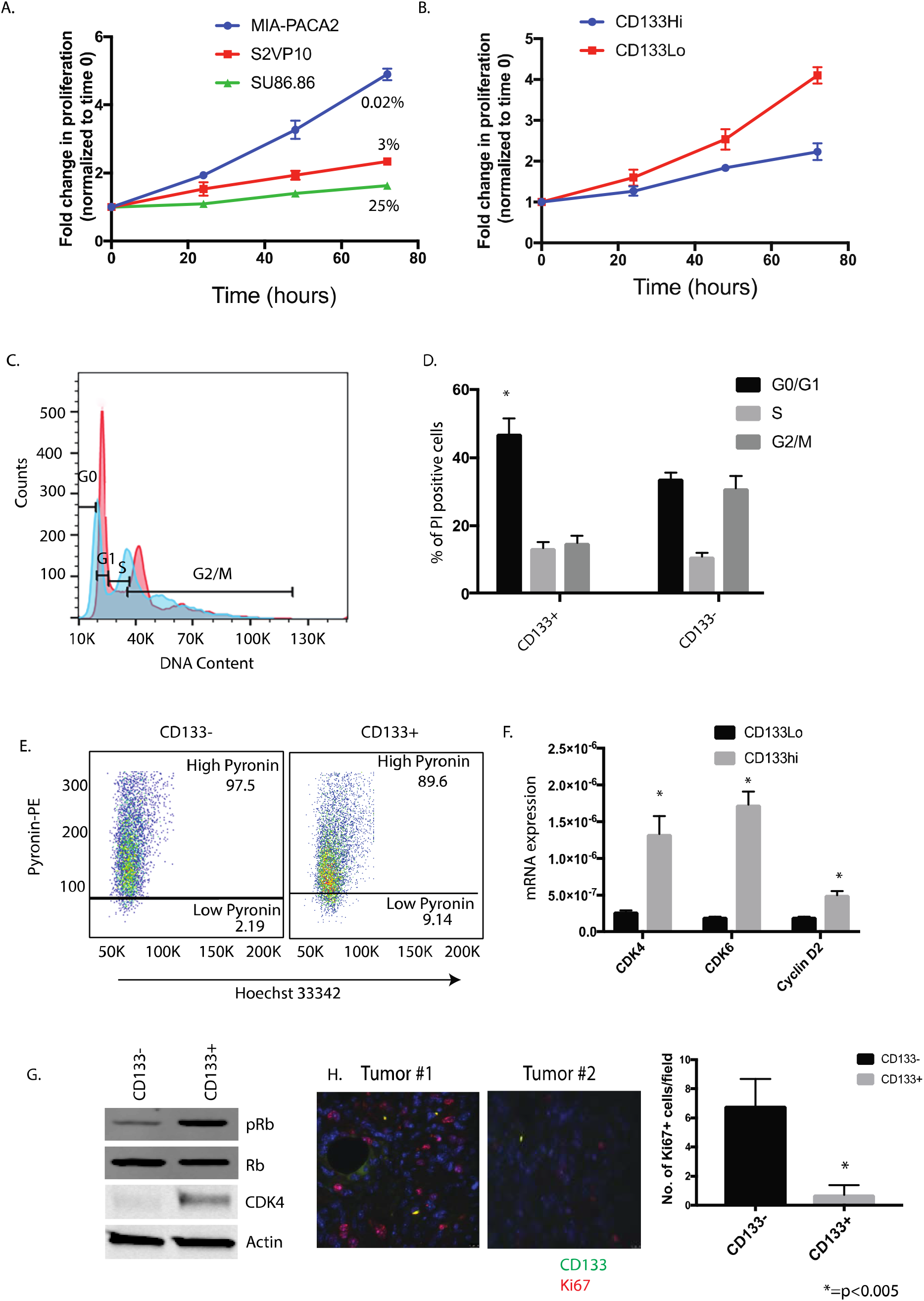
CD133Hi cells are less proliferative and show growth arrest: WST-1 assay was done for cell proliferation. MIA-PACA2 cells with lowest CD133+ population showed highest proliferation while SU86.86 with high CD133+ population showed very little proliferation. Percent of CD133+ population is indicated on the curve (A). CD133 over expressing cells (CD133Hi) also showed increased proliferation compared to CD133Lo cells (B). Cell cycle analysis showed CD133+ cells arrested at G0 state as seen by PI staining. Representative flow analysis (C) and bar graph (D) is shown. CD133+ cells had Low pyronin staining further confirming their G0/G1 arrest (E). Gene expression (F) and protein expression (G) of CDK4/6, Cyclin D2 and phosphorylation of Rb confirmed their arrest in this phase. Immunofluorescence of KPC tumors showed that the CD133+ cells (green) did not stain with Ki67 (red), a proliferative marker, further confirming that these cells were non-proliferative and quiescent (H).

To further confirm these observations *in vivo*, we stained KPC001 derived tumors with CD133 and proliferation marker Ki-67. Our results showed that CD133+ cells typically did not stain with Ki67 (Figure 1H), indicating their quiescence.

### Metabolites in quiescent CD133^Hi^ cells accumulate in biosynthetic pathways

Previously published results from our laboratory showed that CD133+ pancreatic cancer cells had an altered metabolic pathway^26^. To further investigate this, we sorted pancreatic cancer cells isolated from KPC tumors into CD133+ and CD133-populations and labeled them with ^13^C_6_ glucose, and performed a flux analysis on them. Our results showed that in CD133+ cells, the glucose did not follow canonical glycolysis, but instead was routed through the pentose phosphate pathway, to produce lactate as observed by increased 13C labeled glucose (^13^C_6_ Glucose) in the pentose phosphate pathway intermediates (Figure 2A, B). We next validated this by performing an activity assay for glucose 6 phosphate dehydrogenase (G6PD), the first enzyme in the pentose phosphate pathway in the CD133+ and CD133-population. As seen in the flux analysis, CD133+ cells had an increased G6PD activity indicating that in this population glucose was being routed through this metabolic pathway (Figure 2C). To validate if this was a consequence of CD133 expression, we used MIA-CD133^Hi^ cells. As seen with the isolated CD133+ population, MIAPACA2 cells overexpressing CD133 (MIACD133^Hi^) cells showed an increased G6PD activity as well (Figure 2D). Following this, G6PD activity was determined in other pancreatic cancer cell lines with variable expression of CD133. Our results showed that cells with high CD133 had increased G6PD activity compared to those with less CD133 population (Supplementary Figure 2). We analyzed the mRNA expression of the genes involved in the pentose phosphate pathway. As seen in the flux analysis and the enzyme activity assay, the expression of the enzymes in pentose phosphate pathway were increased in CD133^+^ high compared to CD133^−^ cells (Figure 2E). Since pentose phosphate pathway predominantly contributes to the DNA synthesis, we studied the metabolic activity of this pathway in CD133+ cells. We used BrDU, a commonly used thymidine analog that is incorporated into the synthesizing DNA/RNA. Thus, an increase in BrDU incorporation over time would indicate an increase in nucleic acid synthesis. Our study showed that CD133+ cells had almost no BrDU incorporation (Figure 2F), indicating a block in this pathway. This seemed paradoxical as we did not expect quiescent cells to have increased metabolic flux through the biosynthetic pathways like pentose phosphate pathway.

**Figure 2.**
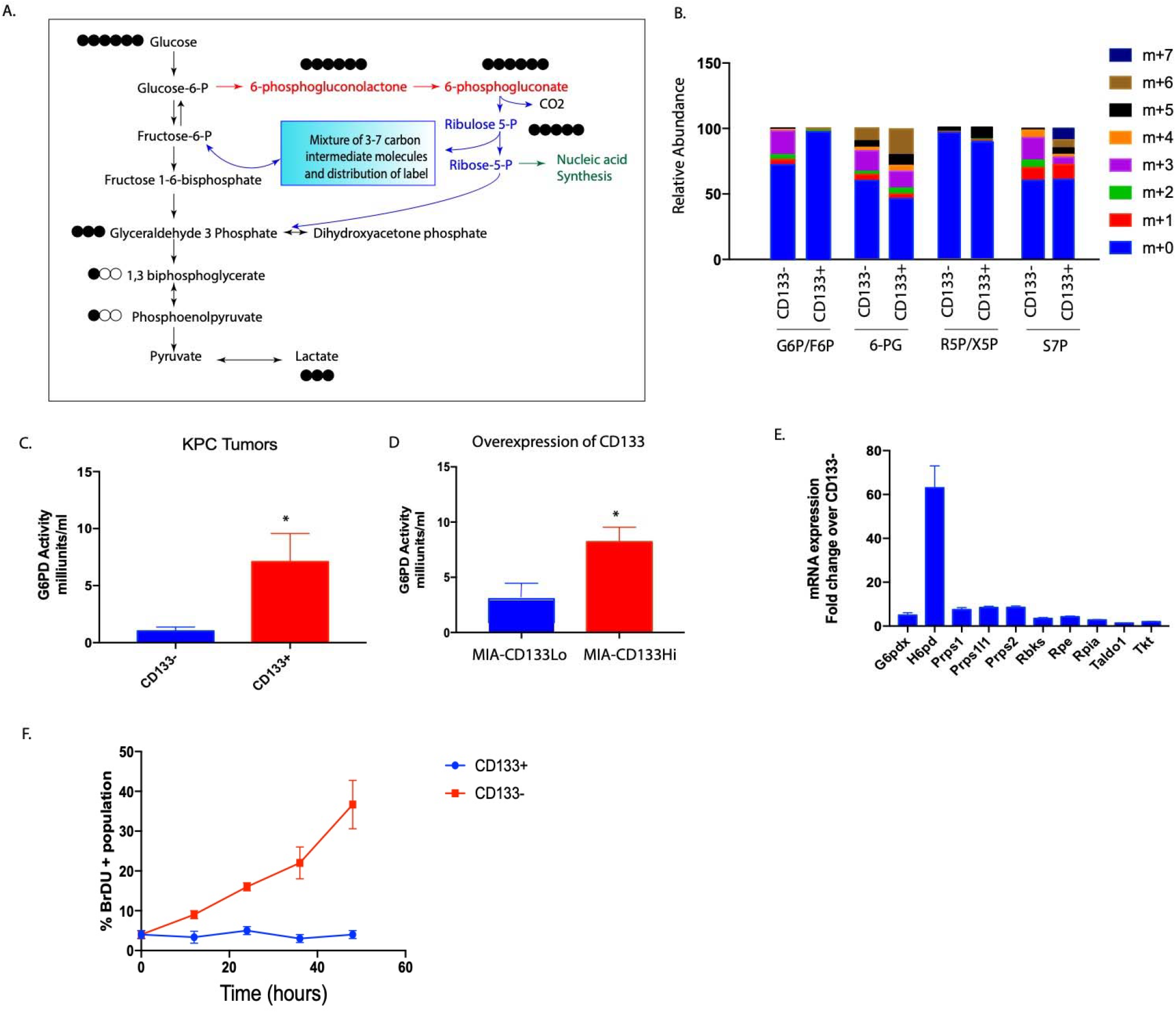
CD133+ cells accumulate metabolites in biosynthetic pathway: ^13^C_6_ glucose labeling of CD133+ cells isolated from KPC tumors showed the glucose was being routed via the pentose phosphate pathway to make lactate instead of following canonical glycolysis (A). Relative abundance of the metabolites (M1-M7) in pentose phosphate pathway in CD133+ and CD133- cells after labeling with ^13^C_6_ glucose for 15 min. Increased accumulation of M+6 isotopologue (6-PG, M+5 isotopologue in R5P/X5P, and M+5, M+6 and M+7 isotopologue can be observed in CD133+ cells (B). CD133+ cells had increased G6PD activity compared to CD133- cells in cells isolated from KPC tumors (C) as well as when CD133 was overexpressed in MIA-PACA2 cell line (D). Similarly, enzymes in the pentose phosphate pathway were overexpressed in CD133+ cells isolated from the KPC tumors (E). Further, CD133+ cells did not incorporate BrDU indicating that they were not synthesizing nucleic acids actively (F).

### 3. Endogenous as well induced expression of CD133 in response to hypoxia, nutritional deprivation and chemotherapy induced stress leads to increased GAS5 expression along with suppression of proliferation

It is known that availability of nutrients regulates cell proliferation by affecting gene transcription. In this context, CD133+ cells that are typically enriched during hypoxia in pancreatic cancer ^21, 25, 26^ are nutritionally deprived. Studies have shown that both hypoxia as well as nutritional deprivation can result in quiescence in a cancer cell population leading to “senescence associated stemness” in a select population of tumor cells. These stressed cells overexpress growth arrest specific 5 or GAS5 gene ^11^. Our analysis showed that cells having increased CD133 expression (whether overexpressed by pCMV-CD133 as in Figure 3A) or endogenous (sorted from a pancreatic tumor) have significantly high GAS5 expression (Figure 3B). Interestingly, when pancreatic cancer cell line MIA-PACA2 was cultured under hypoxia or nutritional stress, GAS5 expression was also increased (Figure 3 C, D). Similar observation was made in other pancreatic cancer cell line as well (Supplementary Figure 3A, B). Previously published results from our laboratory show that under all these conditions, CD133 expression was upregulated as well ^21, 25^. Our previously published work has shown that chemotherapy selected for treatment refractory CD133+ population^25^. To study if GAS5 was upregulated in pancreatic cancer cells under chemotherapy induced stress, we treated KPC001 cells with 100nM Gemcitabine for 7 days and analyzed the GAS5 and CD133 expression. As expected, these cells have extremely high expression of CD133. Along with this, the GAS5 expression was also high in these cells (Figure 3E, Supplementary Figure 3C). We next evaluated if modulation of CD133 affected GAS5 expression. Our results showed that silencing CD133 in S2VP10 cells using siCD133 decreased GAS5 expression (Figure 3F). To study if increased GAS5 expression in CD133Hi cells contributed to its quiescence and growth arrest, we next inhibited GAS5 using siRNA in these cells and studied the effect on proliferation. Our study showed that inhibition of GAS5 increased proliferation in CD133Hi cells as seen by proliferation assays (Figure 3G). To see if these cells now overcame the block in nucleic acid synthesis, we next evaluated their BrDU incorporation. Our study showed that along with increased proliferation, these cells had increased BrDU incorporation after 48h of silencing GAS5 (Figure 3H).

**Figure 3.**
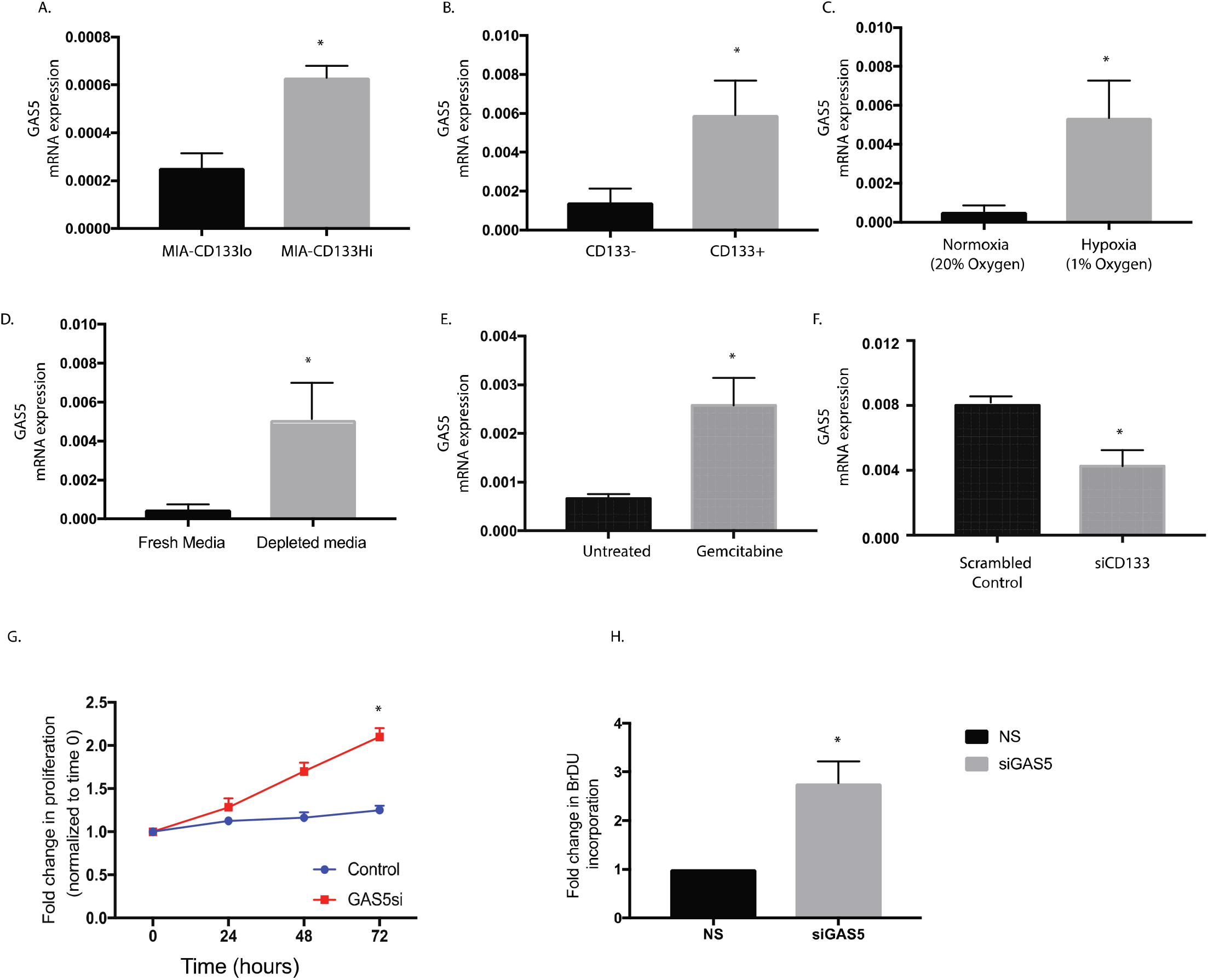
CD133Hi cells have increased GAS5 expression. Pancreatic cancer cells with overexpression of CD133 in MIA-PACA2 cells (A) as well as endogenous CD133+ cells isolated from KPC tumors showed increased GAS5 expression (B). Similarly, exposure to hypoxia (C), nutritional stress (D) and chemotherapy (E) increased GAS5 expression. Silencing CD133 in CD133Hi cells decreased the expression of GAS5 (F). Further, inhibition of GAS5 by siRNA increased proliferation (G) and increased nucleic acid synthesis (H).

### 4. GAS5 is overexpressed in pancreatic tumors

Analysis of public database like Oncomine for GAS5 expression in pancreatic cancer showed that GAS5 was overexpressed in pancreatic tumors when compared to the normal tissues (Figure 4A). Further, GAS5 was observed to be altered in 13% of the samples in the TCGA database hosted at www.cbioportal.org. Among these, mRNA amplification was found to be more prevalent than an alteration in the sequence (Figure 4B). Additionally, GAS5 overexpression was more common in the pancreatic ductal adenocarcinoma and compared to the other types of pancreatic cancers (Figure 4B). To visualize GAS5 in the pancreatic tumors, we next performed a fluorescent in situ hybridization with GAS5 on pancreatic cancer patient tumor slides as well as on implanted tumors with MIA-CD133^Hi^ cells. Our results confirmed that GAS5 was indeed overexpressed in pancreatic cancer (Figure 4C, D). Further, expression of GAS5 increased as the tumor progressed (Figure 4E).

**Figure 4.**
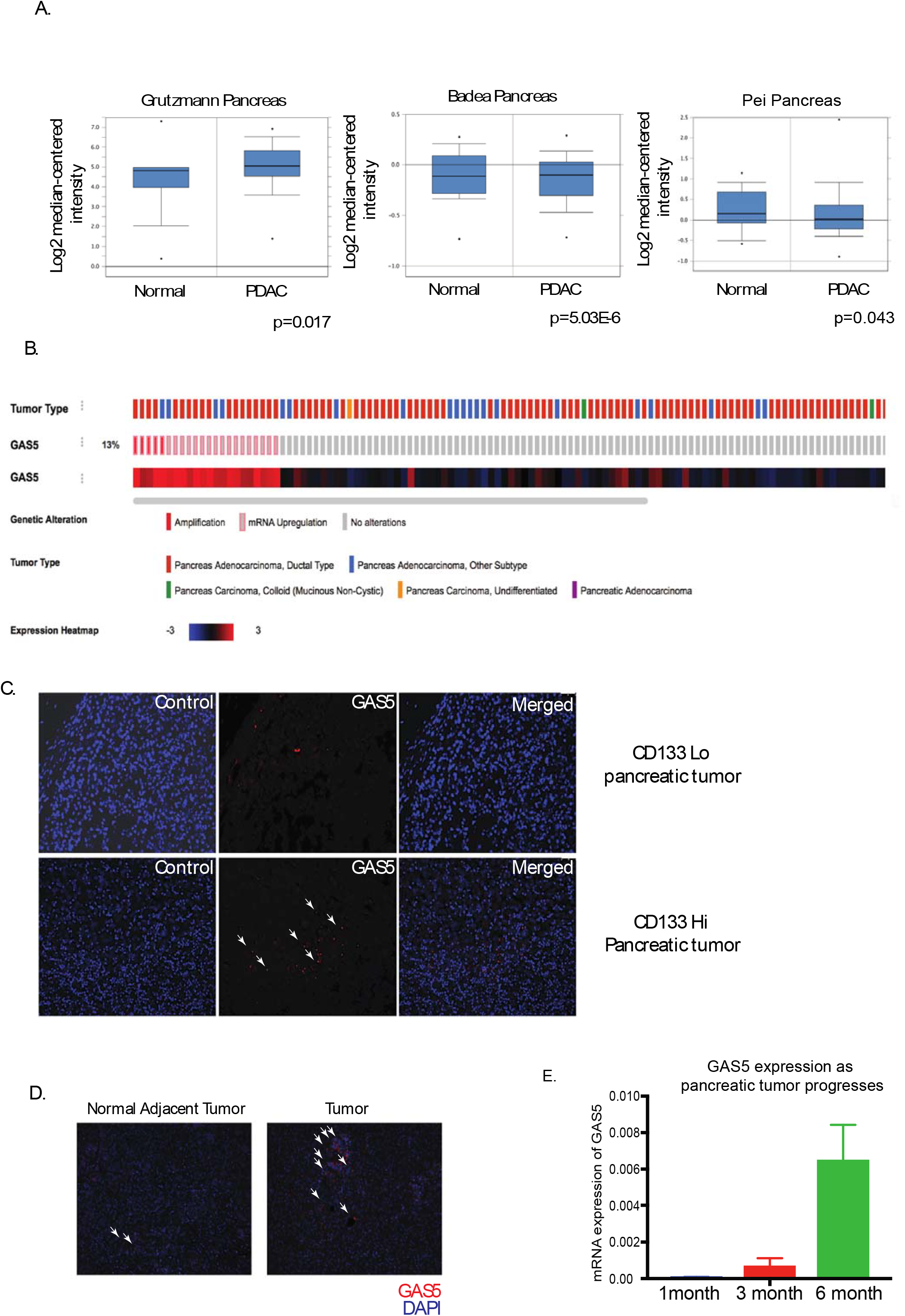
GAS5 was overexpressed in pancreatic tumors. Analysis of Oncomine database showed that GAS5 was increased in pancreatic tumors (A). Further, TCGA database at www.cbioportal.org showed that GAS5 was altered in 13% of the pancreatic tumors and most of the alterations were due to mRNA upregulation and occurred in pancreatic adenocarcinoma (B). FISH confirmed increased GAS5 in CD133hi cells (C) as well as in patient tumors (D). Expression of GAS5 also increased as the tumor progressed in a KPC mouse.

### 5. GAS5 expression and function was regulated by Sox2 in CD133Hi pancreatic cancer cells

We and others have shown that hypoxia increases expression and activity of self-renewal proteins like Sox2/Oct4 and Nanog^21^. Similarly, nutritional deprivation as well as chemotherapy also induced expression of Sox2, Oct4 and Nanog expression ^25^. Literature shows that GAS5 is regulated by Sox2 ^34, 38^. To see if in CD133+ pancreatic cancer cells Sox2 regulated GAS5 mediated growth arrest, we next silenced Sox2 (with siRNA) and studied the expression of GAS5 in CD133^Hi^ cells. As seen in other cancers, inhibition of Sox 2 decreased GAS5 expression (Figure 5A). Additionally, inhibition of Sox2 also decreased expression of CDK4/Cyclin D2 in the cells at the transcriptional (Figure 5 B, C). Since GAS5 was being downregulated with Sox2 inhibition, we hypothesized that this would release the “brake” from the proliferation arrest in the CD133+ cells and promote nucleic acid synthesis. Our study showed that indeed inhibition of Sox2 led to increased proliferation of CD133Hi cells (Figure 5D) as well as increased BrDU incorporation in CD133Hi cells indicating an active nucleic acid synthesis (Figure 5E).

**Figure 5.**
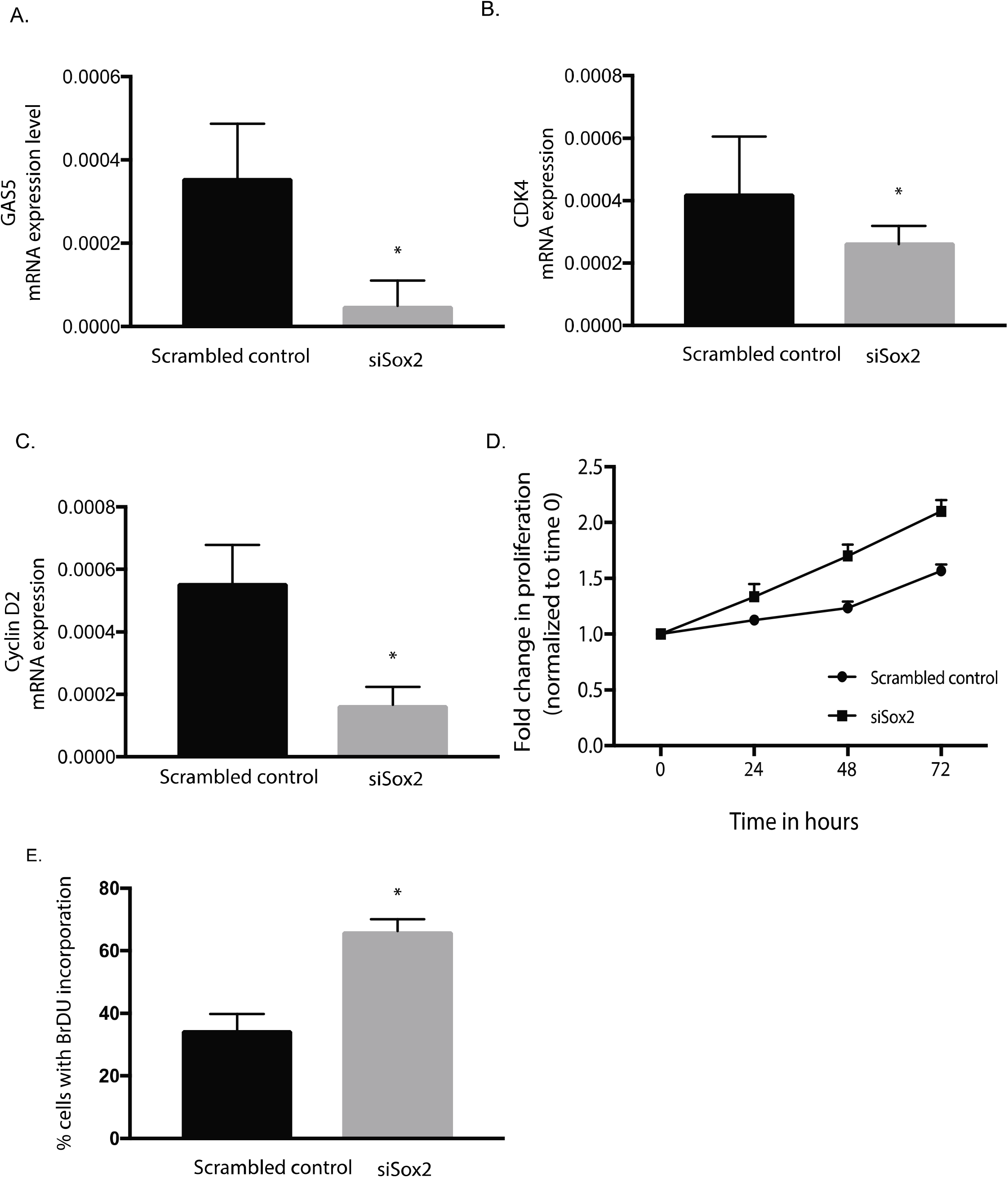
Gas5 expression is regulated by Sox2. Inhibition of Sox2 by siRNA decreased GAS5 mRNA levels (A) along with expression of cell cycle genes CDK4 (B), Cyclin D2 (C) at the transcriptional level. Further, inhibition of Sox2 increased proliferation (D) along with increase in % cells with BrDU incorporation indicating an increase in nucleic acid synthesis (E).

### 6. GAS5 suppressed proliferation and downstream signaling in CD133^Hi^ cells by inhibiting GR mediated signaling

Previous studies have shown that CDK4/ Cyclin D axis can be regulated by glucocorticoid receptors (GR) mediated signaling ^30^. Further, GR transcriptional activity is regulated by GAS5, which binds to the nuclear receptor GR and inhibits its activity. GR is an intracellular ligand dependent transcription factor that binds to GR elements (GRE) in the promoter region of select genes that regulate proliferation. To study if CD133Hi cells with high GAS5 expression consequently had low GR transcriptional activity, we next evaluated GR transcriptional activity using a dual luciferase reporter plasmid. Our study showed that cells with increased GAS5 expression had decreased GR transcriptional activity and inhibition of GAS5 by siRNA increased GR transcriptional activity (Figure 6A). Since Sox2 was responsible for GAS5 transcription, we hypothesized silencing Sox2 will have a similar increase in GR transcriptional activity. Our results confirmed this hypothesis when Sox2 silencing increased GR transcriptional activity (Figure 6A), establishing that Sox2-GAS5 negatively regulated GR activity. Since inhibition of GAS5 suppressed GR activity, we next tested the expression of CDK4/CyclinD expression along with Rb phosphorylation. Our results showed that as expected inhibition of GAS5 decreased expression of CDK4/CyclinD2 expression and decreased Rb phosphorylation (Figure 6 B, C). This was consistent with the decreased proliferation observed with GAS5 inhibition in Figure 3G.

**Figure 6.**
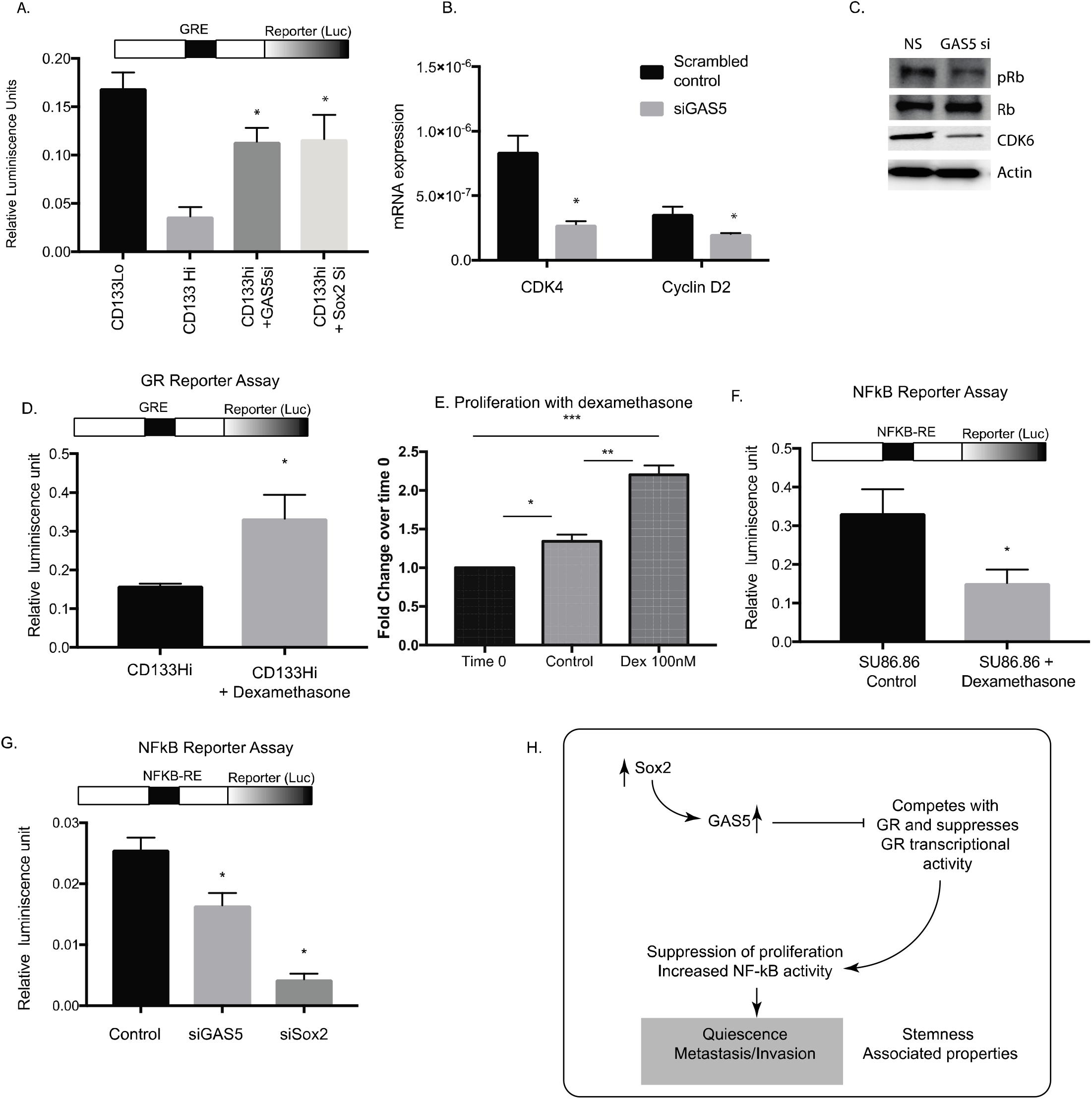
GAS5 regulated Glucocorticoid mediated signaling in PDAC CD133Hi cells. CD133Hi cells had decreased GR activity as seen by dual luciferase reporter activity assay. Silencing GAS5 and Sox2 increased GR transcriptional activity (A). Further, inhibition of GAS5 decreased cell cycle genes like CDK4 and Cyclin D2 at transcriptional (B) and well as translational (C) level. Upon treatment with glucocorticoid, dexamethasone, GR transcriptional activity in CD133Hi cells was increased, indicating a rescue from GAS5 mediated inhibition (D). Proliferation of CD133Hi cells SU86.86 was also increased upon treatment with dexamethasone (E). NF-kB transcriptional activity was negatively regulated by dexamethasone in CD133Hi cells (F). Further, silencing GAS5 and Sox2 inhibited NF-kB transcriptional activity (G). Schematic figure showing how Sox2-GAS5 axis modulates “stemness” associated properties like quiescence in metastasis in CD133hi cells.

To study if the GAS5 mediated growth arrest in CD133+ cells was mediated by GR, we stimulated CD133^Hi^ cells (that typically showed low GR activity) with dexamethasone (a GR agonist). Our results showed that stimulation of GR did not change GAS5 expression in these cells (Supplementary Fig 4A), however, the GR transcriptional activity was significantly increased (Figure 6D). Consistent with this, the cells showed an increase in in proliferation when treated with dexamethasone (Figure 6E). This showed that growth arrest in CD133^hi^ cells was being regulated by GR mediated transcriptional activity.

Previous studies from our lab have shown that CD133+ cells are highly metastatic and this is regulated by the IL1-NF-kB axis^24, 27^. GR has been known to negatively regulate NF-kB ^39^. This seemed consistent with our current observation that CD133Hi cells had increased GAS5 expression, had low GR transcriptional activity and subsequently high NF-kB activity in cells^25^. To study if GR modulation affected NF-kB activity, we next stimulated GR in SU86.86 cells (that have high CD133 and GAS5 expression). We hypothesized that upon stimulation of GR, the inhibition effect of GAS5 on GR will be overcome. Our results showed that upon stimulation of GR (with dexamethasone), NF-kB activity in these cells was decreased (Figure 6F). Our results also showed that silencing Sox2 or GAS5 decreased NF-kB reporter activity, further confirming the role of Sox2-GAS5 axis in mediating function of CD133Hi cells (Figure 6G).

## Discussion

Quiescence has been correlated with “cancer stemness” for a very long time. Early literature in the field reflected that the cancer stem cell (CSC) population was resistant to therapy because of their quiescence, as standard chemotherapy targeted rapidly proliferating cells ^1, 29, 33^. Quiescence properties in a stem cell can be controlled by intrinsic signaling pathways as well as influenced by the tumor microenvironment like hypoxia, interaction with stroma, as well as chemotherapy induced toxicity. Among intrinsic factors, the metabolic property of the cells as well as oncogenic signaling pathways like mTOR signaling play a vital role in determining quiescence ^14^. Recent studies have shown that mitotically quiescent cancer stem cells in solid cancers like breast cancer tend to be more aggressive than the ones that are not mitotically quiescent ^4, 28^. Consistent with this, previous results from our laboratory had identified a quiescent, rare population within pancreatic tumors represented by surface expression of CD133. This population correlated with invasiveness of the pancreatic cancer cells and tumors ^24, 25, 27^ and was found to be responsible for initiating tumors at very low dilutions^2^. Further, results from our lab showed that this CD133+ population was enriched upon stress conditions like nutritional deprivation, hypoxia as well as chemotherapeutic stress^21, 25, 26^, and was identified as a “tumor initiating cell” even though it was not the cell of origin for pancreatic cancer.

Metabolic pathways in cancer have been recognized among the new “hallmarks of cancer”^8^. However, metabolic differences between actively proliferating and non-proliferating cells have not been studied extensively. Quiescent cells are typically known to favor catabolic metabolism compared to the proliferating cells^3, 37^. However, within a tumor not all cells proliferate equally, and thus resulting in a metabolic heterogeneity within the tumor^20^. Interestingly, metabolic flux analysis on isolated CD133+ cells from murine pancreatic tumor cells (KPC001) showed that in spite of being quiescent, a large number of metabolites were being fluxed through biosynthetic pathways like pentose phosphate pathway (Figure 2). This appeared paradoxical, as according to literature, quiescent cells need to be reliant on catabolic rather than biosynthetic pathways in order to maintain homeostasis.

Along with the observation that the CD133+ cells in a tumor had an active pentise phosphate pathway, we also observed that irrespective of whether the cells have endogenously high expression of CD133 or CD133+ population was enriched by microenvironmental cues, these cells always had an overexpression of GAS5 (Figure 4). GAS5 or Growth Arrest Specific 5 factor is a long non-coding RNA that has been reported to promotes cancer cell proliferation by regulating its cell cycle^9, 16, 32^. Functionally, it has been reported that GAS5 lncRNA acts as a “riborepressor” for glucocorticoid receptor^11^. In cells that are arrested for growth, GAS5 is expressed abundantly and competes with glucocorticoids by binding to the DNA binding domain of glucocorticoid receptors (GR) thereby transcriptionally suppressing the GR regulated genes^11^. Our study shows that in pancreatic cancer, CD133+ cells have a high expression of GAS5 (Figure 3), which is regulated by the self-renewal transcription factor Sox2 (Figure 5). Consistent with this, these cells are arrested for growth at the G0/G1 phase (Figure 1). Additionally, these cells have low GR transcriptional activity (Figure 6). Further, inhibition of GAS5 in CD133+ cells, results increased proliferation, and a release from the cell cycle arrest (Figure 4). Furthermore, inhibition of utilization of glucose by 2DG or inhibiting pentose phosphate pathway by DCA has no effect on GAS5 expression (Supplementary Figure 3).

Recent studies have shown that a certain population of cells within the tumor may undergo chemotherapy (or stress) associated quiescence. This helps the population to tide over the unfavorable environment. Upon removal of the stress, these cells reactivate their proliferative phenotype by overcoming their growth arrest^22^. At this stage, the cells undergo a conversion to an aggressive growth phase that is extremely invasive and typically do not respond to therapy. Studies from our laboratory show that CD133+ population is enriched by microenvironmental “stressors” like hypoxia, associated nutritional deficiency as well as chemotherapy. This further increases their GAS5 levels and promotes quiescence. Additionally, GR signaling is known to negatively affect inflammatory pathways by regulating NF-kB. Our study shows that GAS5 mediated suppression of GR activity actually affects NF-kB signaling (Figure 6 G, H).

Thus, our study shows for the first time that in pancreatic cancer, lncRNA GAS5 acted as a molecular rheostat in response to microenvironmental cues, regulating aggressiveness of the CD133+ population. In response to oncogenesis associated stress like hypoxia and nutritional deprivation, a small population of cells, represented by CD133 expression tend to become quiescent by undergoing growth arrest and suppression of proliferation by overexpressing GAS5. GAS5 thus negatively regulated GR transcriptional activity contributing to this growth arrest. This regulation of “growth plasticity” in which metabolites could be stored in biosynthetic pathways during the quiescent phase of the cell cycle and not actively used provides this CD133+ population a survival advantage in which the cells can survive the unfavorable microenvironment by upregulating GAS5. Upon escaping the unfavorable condition, GAS5 expression is decreased and the “brakes” are released allowing CD133+ cells to draw upon the stored metabolites in biosynthetic pathways and jumpstart the processes required for active proliferation resulting in an aggressive tumor.

## Acknowledgement

The authors would like to acknowledge Dr. Charles Burant and Dr. Maureen Kachman Michigan Regional Comprehensive Metabolomics Resource Core, University of Michigan for the metabolic flux analysis. The authors would also like to acknowledge the University of Miami Sylvester Cancer Center Flow Cytometry core for help with the flow cytometry based experiments.

