## Supplementary figures and images for "Long non-coding RNA GAS5 acts as proliferation “brakes” in CD133+ cells responsible for tumor recurrence"

### Supplementary Figure

#### Slide 1
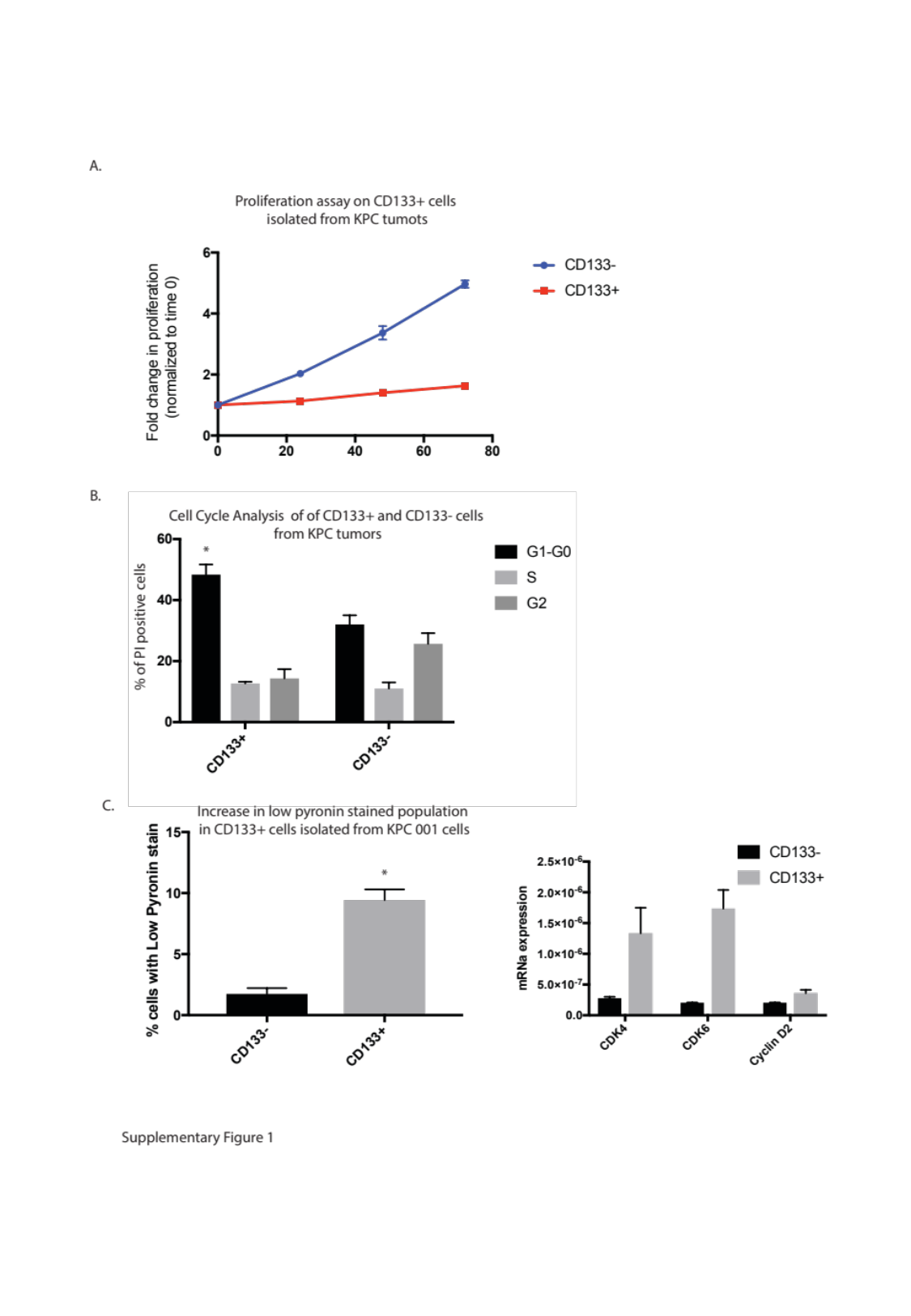

#### Slide 2
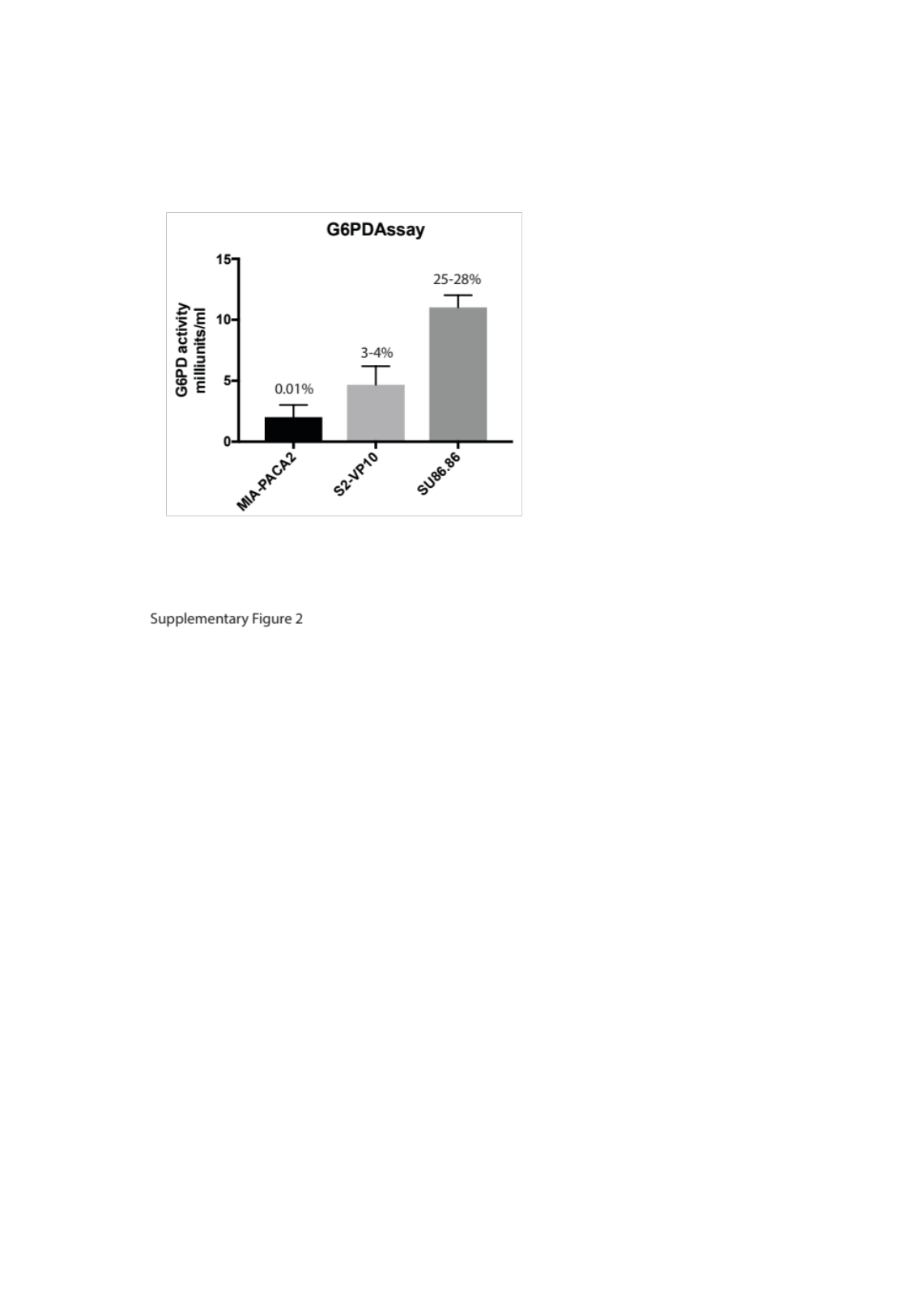

#### Slide 3
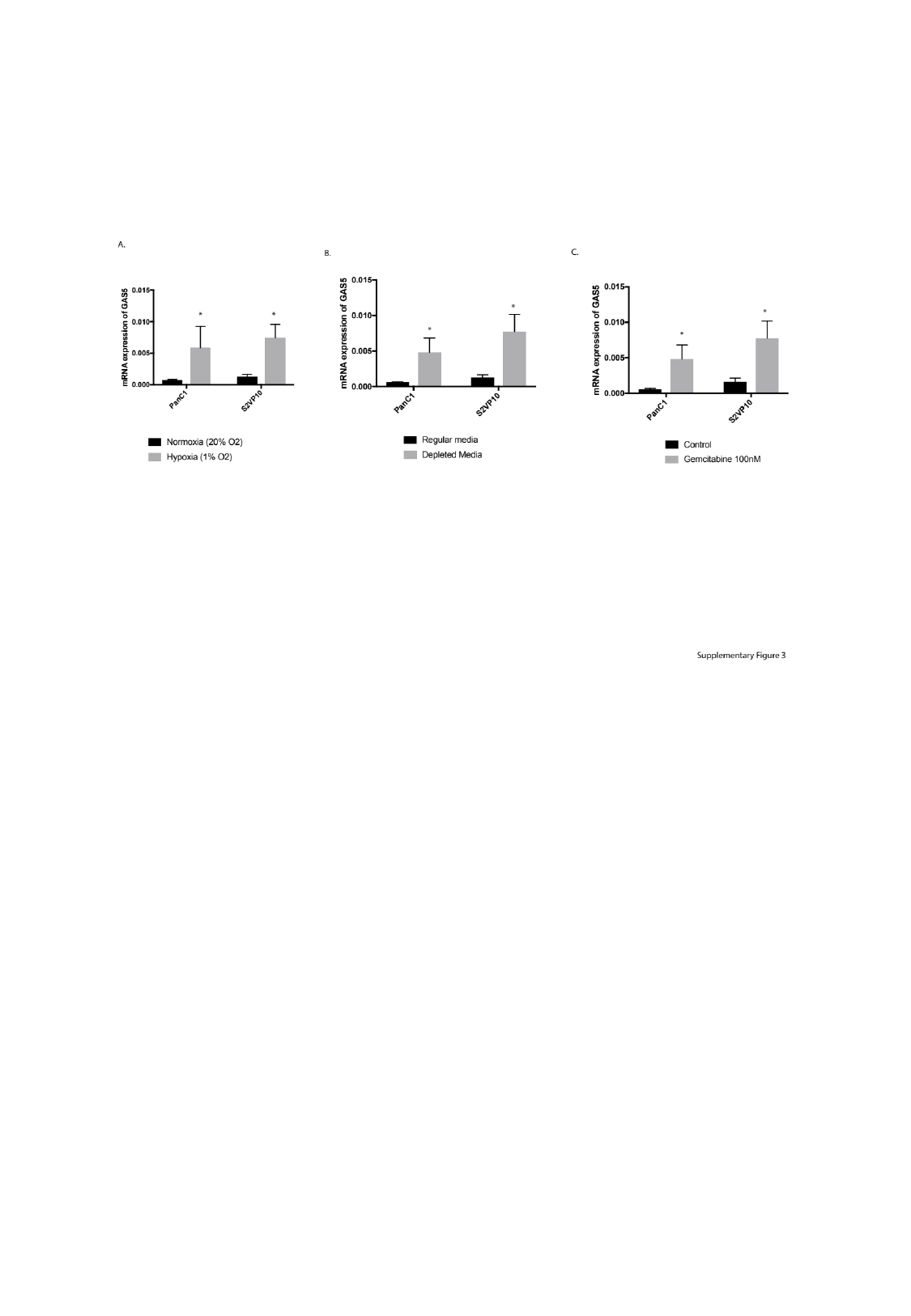

#### Slide 4
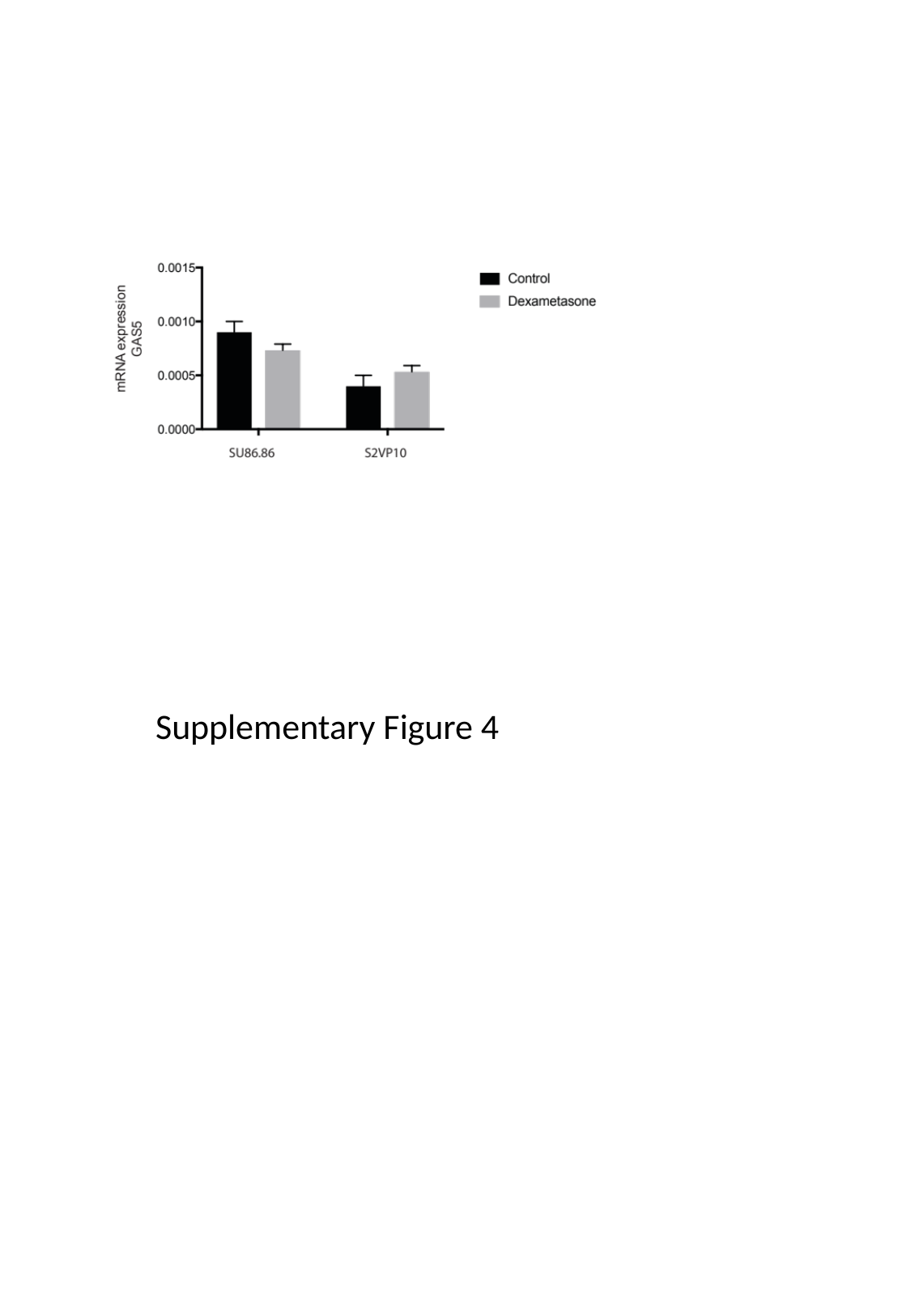

Supplementary Figure 4
